# Highly conserved talin isoforms encode distinct mechanical responses and ligand-binding capabilities

**DOI:** 10.64898/2026.11.05.764451

**Authors:** Yanyu Guo, Durgarao Guttula, Mingxi Yao, Daniel M. Cooper, Neil Ball, Benjamin Klapholz, Alexa Price, Alan Howe, Benjamin T. Goult, Jie Yan

## Abstract

Talin is a central mechanosensitive adaptor that links integrins to the actin cytoskeleton and converts mechanical force into biochemical signals. Vertebrates express two highly homologous talin isoforms, talin-1 and talin-2, yet the molecular basis for their nonredundant functions remains poorly understood. Here, we investigate how isoform-specific sequence differences alter the mechanical response and ligand-binding capabilities of the talin rod. Using single-molecule force spectroscopy and biochemical binding assays, we show that talin-2 preserves the overall mechanical architecture of talin-1 but exhibits distinct mechanical responses in key rod regions. Talin-2 R7-R9 exhibits a previously unobserved three-domain cooperative unfolding pathway, while an alternative sequential pathway reveals early unfolding of R9 at substantially lower force than in talin-1. Strikingly, talin-2 R4-R6 has acquired two additional mechanically regulated interactions: a previously unrecognized vinculin-binding site in R5 and an A-kinase anchoring protein (AKAP) region within R5-R6 that recruits protein kinase A (PKA). Together with the conserved R9 AKAP site, this establishes two spatially separated, mechanically regulated PKA-anchoring regions within the talin-2 scaffold. These findings establish that isoform-specific sequence divergence not only reshapes the mechanical response of the talin scaffold but also expands its repertoire of force-regulated interactions, providing a mechanism for the assembly of different structural and signaling complexes in an isoform-dependent manner.

**Significance Statement:** Cells sense mechanical forces through proteins that change shape under tension, yet how highly conserved mechanosensitive proteins acquire specialized functions remains poorly understood. Talin is a force-bearing scaffold in integrin adhesions, with two closely related isoforms that have nonredundant physiological roles. Here, we show that talin-2 exhibits altered mechanical responses, including a previously unrecognized three-domain cooperative unfolding pathway. Strikingly, talin-2 also contains additional mechanically regulated binding sites for vinculin and protein kinase A, expanding its capacity to recruit structural and signaling proteins in response to force. These findings reveal how sequence divergence between closely related mechanosensitive proteins can generate different mechanical responses and promote the assembly of distinct molecular complexes, providing a mechanism for isoform-specific mechanotransduction.

## Introduction

Cells sense and respond to mechanical forces through specialized protein assemblies that couple adhesion receptors to the cytoskeleton and intracellular signaling pathways (1). Central to this machinery is talin, a large mechanosensitive adaptor that links integrins to actin and converts mechanical load into biochemical signals (2). Talin consists of an N-terminal FERM head domain (F0-F3), which binds integrin cytoplasmic tails and promotes integrin activation (3, 4), and a long C-terminal rod composed of 13 helical bundle domains (R1-R13) (5). Many of these rod domains harbor cryptic ligand-binding sites that become exposed upon mechanical unfolding, allowing talin to function as a force-dependent scaffold that couples mechanical input to ligand recruitment and signaling output (6–9).

Force-dependent signaling by talin arises from the mechanical behavior of individual rod domains that act as binary conformational switches. Under tension, these helical bundles transition between folded and unfolded states, exposing cryptic binding sites that are inaccessible in the folded structure. Because individual domains differ in mechanical stability, the talin rod unfolds in a defined hierarchy, allowing distinct binding sites to become accessible at different force thresholds. Consequently, even subtle changes in the mechanical stability of individual domains could alter how the talin rod responds to force and recruits its binding partners.

Vertebrates express two highly homologous talin isoforms, talin-1 and talin-2, which share approximately 74-76% amino acid identity and an essentially identical domain architecture (10–13). Despite this strong conservation, the two isoforms are not functionally redundant. Talin-1 is ubiquitously expressed and is indispensable for early embryonic development, whereas talin-2 shows a more restricted expression pattern, with particularly high levels in striated muscle and brain, and its loss leads to progressive myopathy rather than embryonic lethality. At the cellular level, talin-1 is enriched in small, nascent adhesions near the cell edge, whereas talin-2 preferentially accumulates in larger, more central and fibrillar adhesions. The functional importance of talin-2 is further highlighted by the identification of a *TLN2* missense variant, S339L, associated with congenital camptodactyly, a condition characterized by permanent flexion contractures of the fingers (14). Sequence divergence is unevenly distributed along the talin rod, with regions such as R5 showing pronounced differences between the isoforms despite strong conservation of their respective sequences across species. Previous cellular studies have demonstrated that talin-1 and talin-2 experience different mechanical tensions and mediate distinct responses to extracellular rigidity, with these differences dependent on the talin rod (15). These differences raise the question of how two proteins with such highly conserved architectures have evolved distinct mechanosensitive functions.

Two important examples of mechanically regulated ligand recruitment by talin are vinculin and protein kinase A (PKA). Vinculin reinforces integrin-actin linkages by binding to amphipathic helices, termed vinculin-binding sites (VBSs), that are buried within folded talin rod domains and exposed upon mechanical unfolding (7, 16–18). Vinculin binding stabilizes unfolded talin states and promotes adhesion reinforcement, providing a direct link between mechanical force and structural stabilization. In addition to recruiting structural proteins, talin also functions as a mechanically gated A-kinase anchoring protein (AKAP), binding PKA through a cryptic site exposed by mechanical unfolding (19). Together, these interactions illustrate how the mechanical unfolding hierarchy of the talin rod determines when and where structural and signaling proteins can be recruited.

Here, we investigate how sequence divergence between talin-1 and talin-2 influences their mechanical responses and force-dependent ligand interactions. Using single-molecule force spectroscopy and biochemical binding assays, we show that the two isoforms preserve a common mechanical architecture but exhibit distinct unfolding behaviors in specific rod regions. Talin-2 R7-R9 displays striking interdomain cooperativity and an altered unfolding hierarchy, while differences in the R4-R6 region are accompanied by previously unrecognized ligand-binding interactions, including a vinculin-binding site in R5 and an additional PKA-anchoring region within R5-R6. Structural comparisons identify isoform-specific buried polar residues that provide a potential explanation for the altered mechanical stability of these conserved domains. Together, these findings demonstrate how highly conserved talin isoforms encode distinct mechanical responses and ligand-binding capabilities, revealing how sequence divergence can diversify the mechanosensitive functions of a conserved protein scaffold.

## Results

### Mechanical unfolding of talin-2 rod domains

Talin-1 and talin-2 share a conserved rod architecture comprising 13 helical bundle domains (R1– R13), but exhibit substantial sequence divergence across individual domains (Fig. 1A). Our previous single-molecule studies showed that talin-1 rod domains unfold sequentially under force in a defined mechanical hierarchy that regulates the exposure of cryptic ligand-binding sites (8). Although the talin rod contains thirteen domains, it exhibits up to twelve discrete unfolding steps. This is because R8 is inserted within R7 (20) and is not directly loaded under tension until R7 unfolds, resulting in the cooperative unfolding of the R7–R8 module (8).

To determine whether the global mechanical architecture of the talin rod is conserved between isoforms, we performed repeated force-ramp measurements on individual talin-2 molecules at a constant loading rate of 2 pN/s. In each cycle, the talin-2 rod was stretched from 1 to approximately 30 pN and subsequently relaxed to below 2 pN to allow unfolded domains to refold, enabling multiple force-extension traces to be recorded from a single tether.

Representative force-bead height traces from repeated stretching of a single talin-2 rod are shown in Figure 1B. Each trace displays a series of discrete extension steps corresponding to the unfolding of individual helical bundle domains. Up to twelve unfolding events were observed per stretch, consistent with the thirteen-domain architecture of the talin rod and the mechanically coupled unfolding of R7–R8, as previously observed for talin-1. The overall unfolding behavior of the talin-2 rod is broadly similar to that of talin-1, prompting us to investigate whether individual rod segments exhibit distinct mechanical properties.

**Figure 1.**
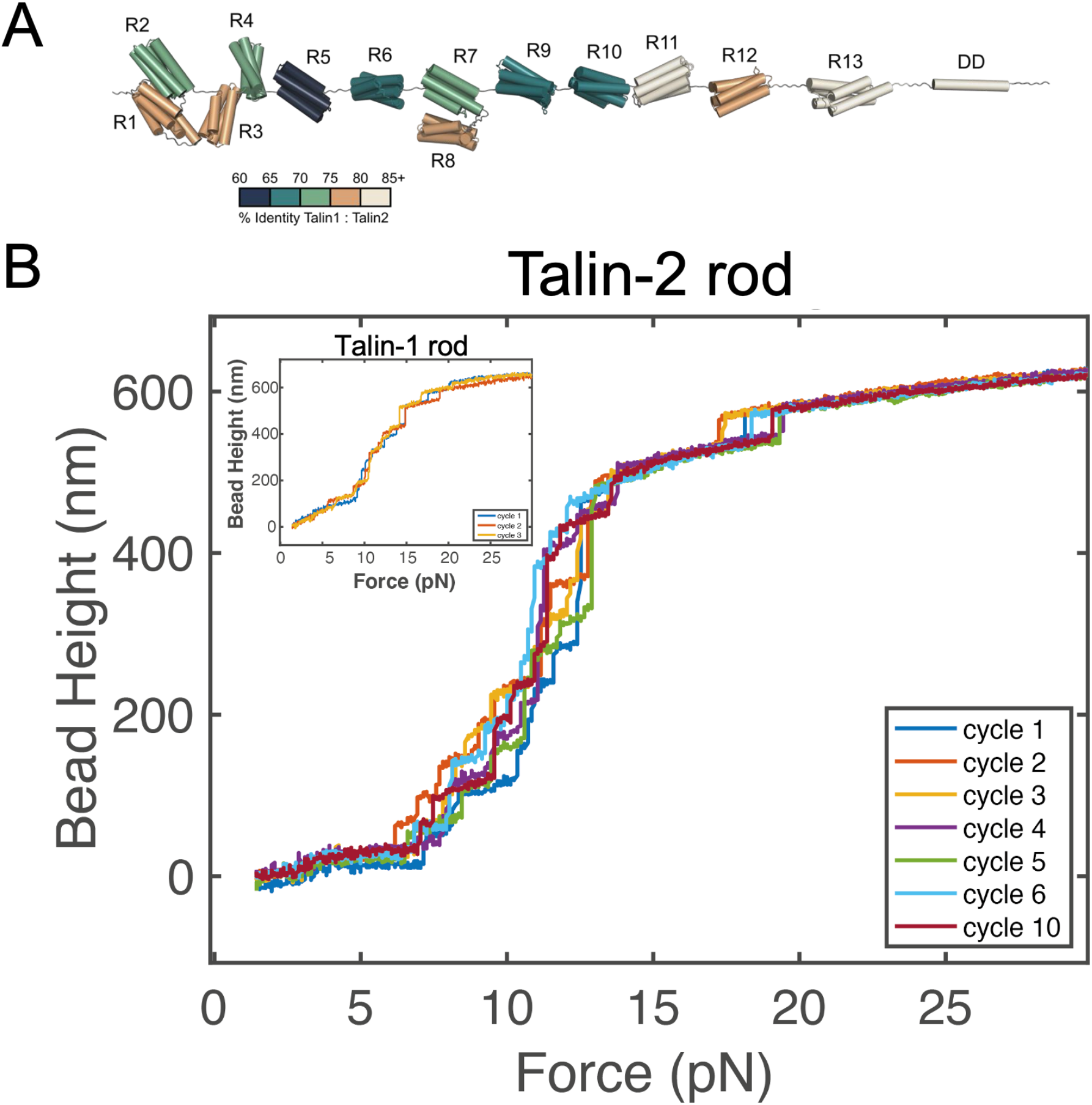
Structural organization and mechanical unfolding of the talin-2 rod. **(A)** Structural representation of the talin rod showing the arrangement of the 13 helical bundle domains (R1-R13) and the C-terminal dimerization domain (DD). Individual domains are colored according to their percentage sequence identity between talin-1 and talin-2, highlighting the nonuniform distribution of sequence conservation along the rod. **(B)** Stretching the talin-2 rod at a loading rate of 2 pN/s produces up to twelve discrete unfolding steps over a force range of 2-30 pN. These stepwise extension events correspond to the unfolding of helical bundle domains and closely resemble the overall unfolding behavior previously observed for talin-1 (inset).

### Isoform-specific retuning of talin rod mechanics

To identify isoform-dependent differences in local mechanical properties, we applied the same repetitive force-cycling protocol to individual talin-2 rod subsegments. This approach yielded hundreds of unfolding events per construct across multiple independent tethers, enabling statistical analysis of unfolding forces and contour-length changes (Fig. 2A,B).

#### R1-R3

The first three domains of the talin-2 rod exhibited two major unfolding transitions indicated by stepwise bead height increments, together with pronounced low-force fluctuations near ∼5 pN (Fig. 2A). These fluctuations are consistent with rapid unfolding-refolding transitions of R3, which occur at approximately 4 pN in talin-2 compared with approximately 5 pN in talin-1 (Fig. 2B), indicating a modest reduction in R3 mechanical stability. Compared with talin-1, the higher-force unfolding transitions in this region are redistributed, with one population shifted toward lower forces, peaking at approximately 12 pN, while another extends to higher forces in talin-2 (Fig. 2C). Together, these changes indicate that the mechanical stability of the R1-R3 segment is selectively retuned in talin-2, with some domains unfolding at lower forces while others exhibit increased mechanical stability.

**Figure 2.**
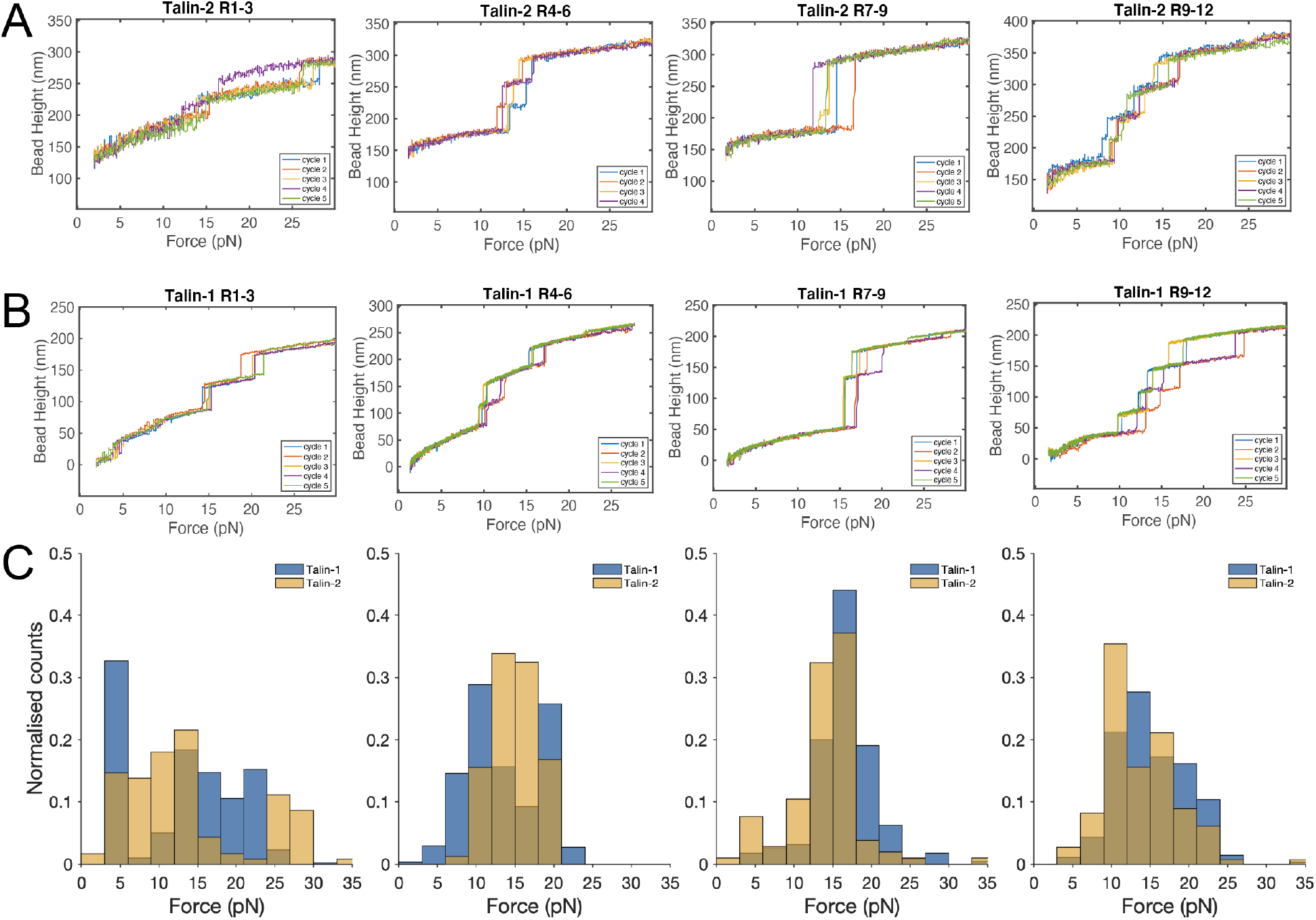
Isoform-specific differences in unfolding behavior across the talin rod. **(A) Talin-2 rod subsegments**. Representative force-bead height traces for R1-R3, R4-R6, R7-R9, and R9-R12 recorded under a constant loading rate. Stepwise increases in bead height correspond to unfolding events. **(B) Talin-1 rod subsegments**. Corresponding force-bead height traces for talin-1 subsegments recorded under identical conditions. **(C) Unfolding force distributions**. Histograms of unfolding forces obtained from repeated measurements of each subsegment (talin-1, blue; talin-2, yellow, overlapping brown) revealing isoform-specific differences in mechanical stability and unfolding behavior.

#### R4-R6

The talin-2 R4-R6 fragment exhibited three unfolding steps clustered within a relatively narrow force range of approximately 10-15 pN (Fig. 2A). In contrast, talin-1 R4-R6 exhibited a broader distribution extending to higher forces (Fig. 2B,C), indicating that the high-force unfolding transition observed in talin-1 is absent from talin-2. Structural analysis of R5 reveals a cluster of threonine residues within the hydrophobic core of talin-2 that is absent from the corresponding domain of talin-1. This arrangement resembles the destabilizing threonine belt previously characterized in talin-1 R3, suggesting that altered hydrophobic core packing may reduce the mechanical stability of talin-2 R5 and account for the loss of the high-force unfolding transition observed in talin-1 R4-R6.

#### R9-R12

The talin-2 R9-R12 fragment exhibited four unfolding steps spanning a broad force range of approximately 10-25 pN (Fig. 2A). Although the unfolding force distributions broadly overlap between the isoforms, talin-1 occasionally exhibited a higher-force unfolding transition that is less frequently observed in talin-2 (Fig. 2B,C), suggesting subtle differences in mechanical stability within this region. However, with the exception of R9, the unfolding steps within R9-R12 cannot be unambiguously assigned to individual domains, and subtle differences in their unfolding order may therefore remain unresolved.

Taken together, these results reveal a redistribution of mechanical stability along the talin-2 rod consistent with targeted tuning of key mechanosensitive segments.

### Talin-2 R7-R9 exhibits a distinct cooperative unfolding pathway and early R9 unfolding

The talin-2 R7-R9 segment exhibited strikingly different mechanical behavior compared with talin-1. Representative force-bead height traces revealed two dominant unfolding pathways (Fig. 3A). In the first pathway, unfolding proceeded in two steps, with an initial ∼40 nm extension corresponding to R9 unfolding, followed by a larger ∼80 nm extension associated with co-unfolding of R7-R8. In the second pathway, a single large ∼120 nm step was observed, indicating cooperative unfolding of all three domains in a single transition. Both pathways were observed across repeated stretching cycles of the same molecule and reproduced across multiple independent molecular tethers, indicating that the distinct unfolding behaviors are intrinsic to the mechanical properties of the talin-2 R7-R9 module rather than arising from differences between individual molecules. In contrast, previous single-molecule studies of talin-1 R7-R9 revealed a sequential unfolding pathway in which the mechanically coupled R7-R8 domains unfold together, followed by a separate R9 unfolding event (8). Notably, the fully cooperative ∼120 nm unfolding transition observed in talin-2 was absent from the talin-1 unfolding pathway.

Analysis of unfolding events in contour length-force space further resolves these pathways into distinct populations (Fig. 3B). A low-force population below ∼10 pN, with contour-length increments of approximately 40 nm, corresponds to early unfolding of R9. A second population at intermediate forces with ∼80 nm increments is associated with co-unfolding of R7-R8. A third population with ∼120 nm increments spans a broad force range and corresponds to fully cooperative unfolding of the R7-R9 tri-domain unit. The substantial overlap in force between these populations indicates that sequential and cooperative pathways coexist over similar force ranges.

Structural studies of the talin-1 R7-R9 module revealed interactions between R8 and R9 in the crystal lattice (21), raising the possibility that analogous interdomain contacts could influence the mechanical behavior of this region. Isoform-specific differences in these interactions may therefore contribute to the altered mechanical coupling observed in talin-2 R7-R9.

Together, our results demonstrate that isoform-specific sequence divergence selectively retunes the mechanical hierarchy of the talin rod, potentially altering the force-dependent exposure of cryptic ligand-binding sites.

**Figure 3.**
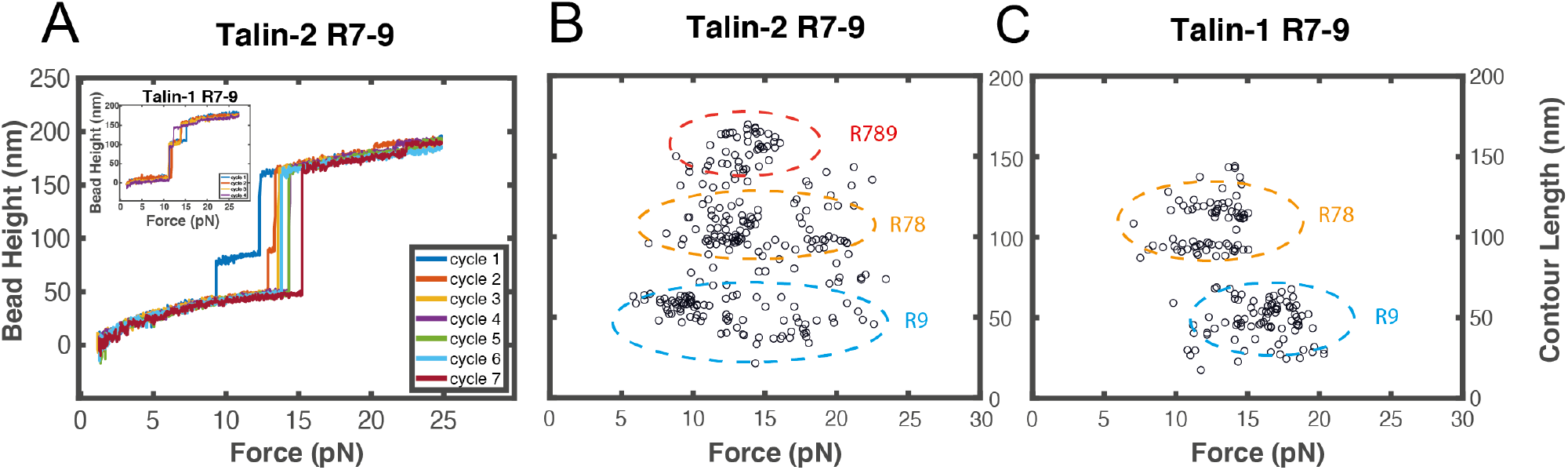
Comparison of the mechanical unfolding behavior of talin-1 and talin-2 R7-R9 modules. **(A)** Representative force-bead height traces of talin-2 R7-R9 recorded during repeated force cycles, with corresponding talin-1 R7-R9 traces shown in the inset. Talin-2 exhibits two distinct unfolding pathways, with R9 unfolding before the cooperative unfolding of R7-R8, or all three domains unfolding cooperatively in a single transition. In contrast, talin-1 exhibits a consistent unfolding hierarchy in which cooperative unfolding of R7-R8 precedes the smaller R9 unfolding step. **(B)** Contour-length changes plotted against unfolding force for talin-2 R7-R9, revealing three populations corresponding to R9 unfolding (∼50 nm contour length change), cooperative R7-R8 unfolding (∼110 nm), and cooperative unfolding of the complete R7-R9 module (∼170 nm). The contour length changes are converted from unfolding step sizes using a bending persistence of 0.8 nm of polypeptide. **(C)** Corresponding contour-length changes plotted against unfolding force for talin-1 R7-R9, showing distinct populations associated with cooperative R7-R8 unfolding and subsequent R9 unfolding.

### Force-dependent vinculin binding reveals an additional high-affinity VBS in talin-2

We next examined how the altered mechanical properties of talin-2 influence force-dependent vinculin recruitment. Single-molecule stretching experiments were performed on talin-2 rod subsegments (R1-R3, R4-R6, R7-R9, and R9-R12) in the presence or absence of 100 nM vinculin head domain (Vd1) (Fig. 4). In the absence of Vd1, all constructs exhibited the expected stepwise unfolding transitions corresponding to domain opening events.

Upon addition of Vd1, the force-bead height traces changed markedly, with discrete unfolding steps progressively reduced or lost over successive force cycles and replaced by smoother extension profiles in all four constructs (Fig. 4A-D, left). This behavior is consistent with Vd1 binding to vinculin-binding sites (VBSs) exposed by mechanical unfolding, thereby stabilizing unfolded states and inhibiting refolding during relaxation to low force. Thus, as in talin-1, vinculin binding to talin-2 remains force-gated.

#### R1-R3

The effects of Vd1 were particularly pronounced in the talin-2 R1-R3 segment. Following Vd1 addition, the number of domains undergoing refolding rapidly decreased to zero over successive force cycles (Fig. 4A), indicating that all three domains contain high-affinity VBSs capable of stably engaging Vd1 and preventing refolding at low force. The altered mechanical stability of talin-2 R1-R3 suggests that these VBSs may become accessible over a broader range of mechanical forces than in talin-1, potentially modifying the force-dependent recruitment of vinculin.

#### R4-R6

The talin-2 R4-R6 segment also showed strong suppression of refolding following Vd1 addition (Fig. 4B), with the number of refolding domains progressively decreasing from three to one. This contrasts markedly with talin-1 R4-R6, in which only R6 has been shown to contain a high-affinity VBS capable of preventing refolding at low force. The suppression of two refolding transitions indicates the presence of an additional high-affinity VBS within talin-2 R4-R6, while the third domain remains able to refold in the presence of Vd1.

#### R7-R9

The talin-2 R7-R9 segment retained one refolding domain even after repeated exposure to Vd1 (Fig. 4C), consistent with R9 lacking a VBS, as previously established for talin-1.

#### R9-R12

Likewise, in talin-2 R9-R12, the number of domains able to refold decreased from four to two after Vd1 addition (Fig. 4D), consistent with the presence of two high-affinity VBS-containing domains in this region, as previously observed in talin-1.

**Figure 4.**
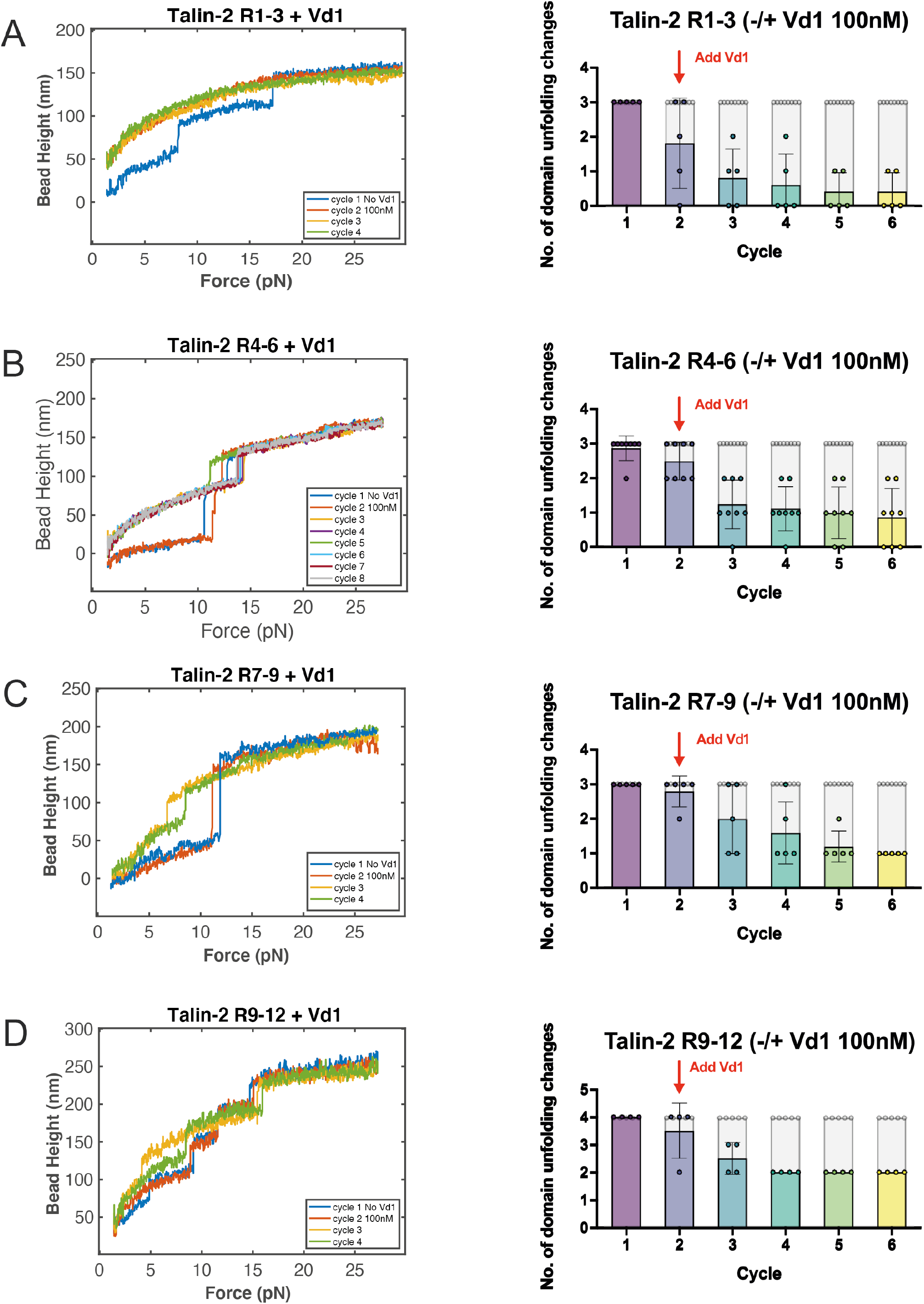
Force-gated vinculin binding reveals an additional high-affinity VBS within talin-2 R4-R6. (**A-D**) Representative force-bead height traces (left) and quantification of the number of domains able to refold during successive force cycles (right) for talin-2 rod subsegments R1-R3 (A), R4-R6 (B), R7-R9 (C), and R9-R12 (D), measured in the absence and presence of 100 nM vinculin head domain (Vd1). Each dot represents an independent tether. The first stretching cycle was recorded in the absence of Vd1, which was then added before subsequent cycles, as indicated. In the presence of Vd1, binding to vinculin-binding sites (VBSs) exposed by mechanical unfolding prevents refolding of the corresponding domains at low force, resulting in a progressive reduction in the number of unfolding steps in subsequent cycles. The extent of refolding suppression differs across subsegments, with R4-R6 showing suppression of refolding in two of its three domains, consistent with an additional high-affinity VBS within this region. All measurements were performed at a loading rate of 2 pN/s in the presence of 2 mM TCEP.

### Biochemical identification of a novel vinculin-binding site in talin-2 R5

The single-molecule experiments revealed an additional high-affinity VBS within talin-2 R4-R6, but did not establish which domain contained the site. To identify the responsible domain, we investigated the ability of isolated talin-2 R4, R5, and R6 domains to form complexes with the vinculin head domain (Vd1) using analytical size-exclusion chromatography (SEC). Each domain was incubated with Vd1 at a 1:1 molar ratio for 16 h at 42 °C to promote transient domain opening and maximize the detection of interactions with cryptic VBSs, as previously described (22) (Fig. 5).

As expected, R6 formed a stable complex with Vd1, with more than 95% of Vd1 incorporated into the complex, while R4 showed no detectable binding, consistent with the corresponding domains in talin-1. Strikingly, R5 also formed a complex with Vd1, identifying a previously unrecognized vinculin-binding site within this domain and confirming that the additional VBS detected in the single-molecule experiments resides in talin-2 R5.

Together, these biochemical results identify R5 as the source of the additional VBS detected in the single-molecule experiments, establishing an isoform-specific expansion of the vinculin-binding repertoire within the talin-2 rod.

**Figure 5.**
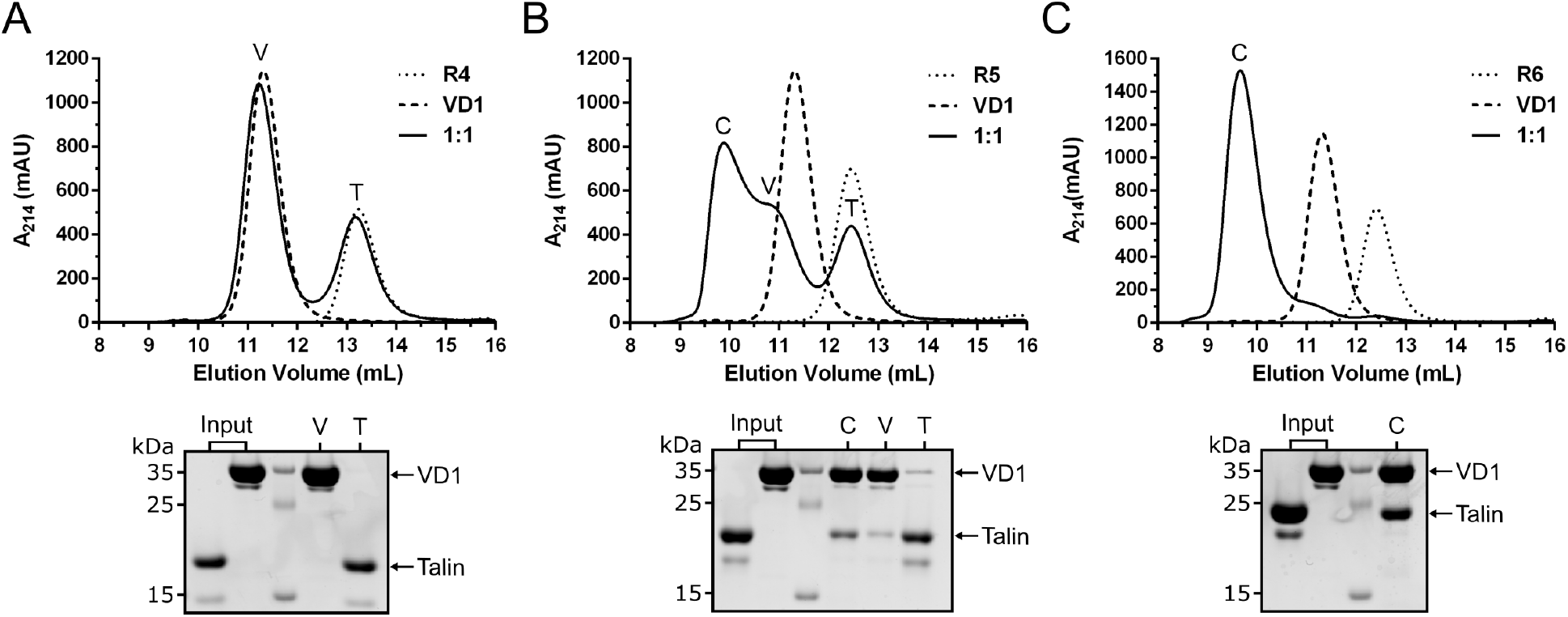
Identification of a novel vinculin-binding site in talin-2 R5. Analytical size-exclusion chromatography (SEC) elution profiles of talin-2 R4 **(A)**, R5 **(B)**, and R6 **(C)** following incubation with vinculin head domain (Vd1) at a 1:1 molar ratio for 16 h at 42 °C. Complex formation is observed for R5 and R6, but not R4, identifying an additional vinculin-binding site in talin-2 R5. Fractions corresponding to the indicated elution peaks were analyzed by SDS-PAGE, with proteins visualized by Coomassie Brilliant Blue R-250 staining (bottom).

### Talin-2 contains two mechanically gated PKA-binding regions at distinct positions along the rod

Having identified an additional vinculin-binding site in talin-2, we next investigated whether isoform-specific divergence had similarly altered the distribution of A-kinase anchoring protein (AKAP) sites along the talin rod. Our previous study established that talin-1 contains a mechanically gated AKAP helix within the R9 region that recruits protein kinase A (PKA) upon mechanical unfolding (19). We examined force-dependent recruitment of PKA using the same repeated force-cycling assay. Talin-2 rod subsegments (R1-R3, R4-R6, R7-R9, and R9-R12) were stretched in the absence and presence of 1 µM PKA, and changes in domain refolding behavior were monitored over successive force cycles (Fig. 6).

The R1-R3 segment showed little or no detectable response to PKA, with unfolding traces remaining largely unchanged over repeated cycles and the number of domains able to refold preserved (Fig. 6A). In contrast, clear effects of PKA were observed in R4-R6, R7-R9, and R9-R12. Following PKA addition, the number of domains able to refold decreased (Fig. 6B-D), consistent with PKA binding to mechanically exposed AKAP helices and preventing refolding at low force.

The suppression of refolding in R4-R6 identifies a previously unrecognized mechanically gated AKAP region in talin-2. The overlapping R7-R9 and R9-R12 segments both contain R9, consistent with the presence of the previously characterized R9-associated AKAP site in both talin isoforms. The additional PKA-binding region in talin-2 R4-R6 therefore expands the kinase-anchoring capacity of the rod, providing a second site through which mechanical forces can regulate the recruitment and spatial organization of PKA.

**Figure 6.**
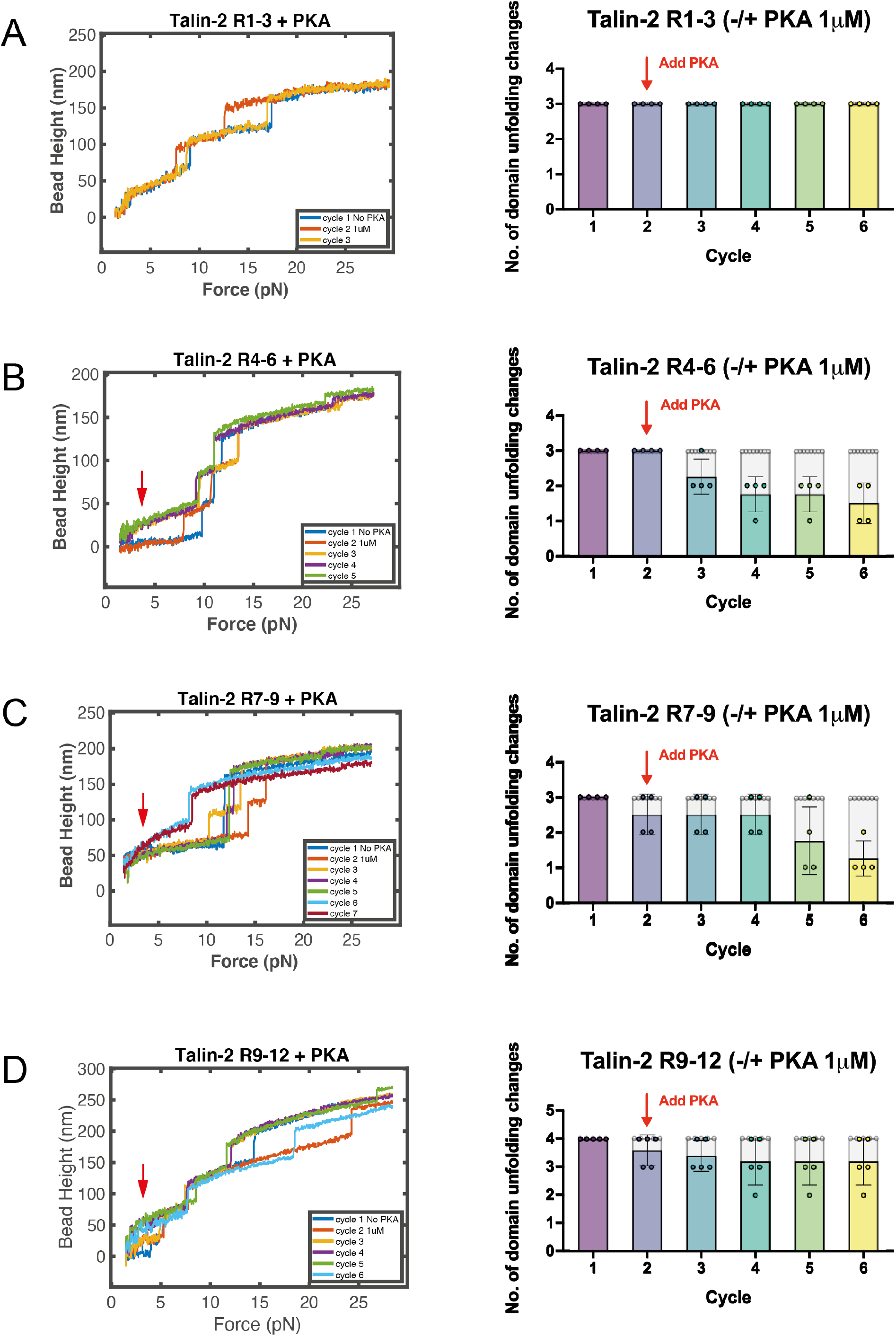
PKA binding reveals two mechanically gated kinase-anchoring regions in the talin-2 rod. (A-D) Representative force-bead height traces (left) and quantification of the number of domains able to refold during repeated force cycles (right) for talin-2 rod subsegments R1-R3 (A), R4-R6 (B), R7-R9 (C), and R9-R12 (D), measured in the absence and presence of 1 µM PKA. Each dot represents an independent tether. The first stretching cycle was recorded without PKA, followed by addition of PKA prior to subsequent cycles, as indicated. In the presence of PKA, R4-R6, R7-R9, and R9-R12 show suppression of one refolding transition, whereas R1-R3 remains largely unaffected. As R7-R9 and R9-R12 overlap at R9, these results are consistent with two mechanically gated PKA-binding regions within talin-2, located in R4-R6 and the R9-containing region. All measurements were performed at a loading rate of 2 pN/s in the presence of 2 mM TCEP.

### Talin-2 R5-R6 contains an additional AKAP region

The single-molecule experiments identified an additional mechanically gated PKA-binding region within talin-2 R4-R6. To investigate this interaction further, we examined the binding of the PKA type II regulatory subunit RIIα to recombinant talin-1 and talin-2 rod fragments using an RIIα overlay assay (Fig. 7). PKA regulatory subunits mediate kinase localization through interactions with amphipathic helices within AKAPs. Interestingly, R4-R6 is among the least conserved regions of the talin rod, with sequence identity between talin-1 and talin-2 falling to 60% in R5, compared with 73% in R4 and 68% in R6 (Fig. 7A).

Using recombinant talin-1 and talin-2 fragments spanning R4-R8 and R9-R10, we observed striking isoform-specific differences in RIIα binding. Both talin-1 and talin-2 R9-R10 bound RIIα, consistent with conservation of the previously characterized AKAP interaction. However, the talin-2 R9-R10 fragment produced a substantially stronger overlay signal than the corresponding talin-1 fragment, despite comparable protein loading as confirmed by Ponceau staining, suggesting enhanced RIIα binding at this conserved site. Talin-2 R4-R8 also showed strong RIIα binding, whereas the corresponding talin-1 fragment showed no detectable interaction (Fig. 7B). These interactions were abolished by the AKAP-competitive peptide Ht31 but not by the inactive control Ht31P, confirming that they involve canonical AKAP-binding interfaces. Thus, talin-2 contains an additional PKA-binding region and exhibits enhanced RIIα binding within the conserved R9-R10 region.

To localize this additional interaction more precisely, we examined binding of the RIIα dimerization/docking (D/D) domain to talin-2 R4-R5 and R5-R6 using NMR spectroscopy. Previous studies of talin-1 R9-R10 demonstrated selective broadening of R9 resonances upon addition of RIIα D/D, while R10 remained largely unaffected, consistent with PKA binding destabilizing the folded R9 domain. Similarly, addition of unlabeled RIIα D/D to talin-2 R5-R6 caused pronounced spectral perturbations, including peak broadening, at a 1:1 molar ratio (Fig. 7D). In contrast, R4-R5 showed little detectable change even at a 1:3 molar ratio (Fig. 7C). These results localize the additional PKA-binding interaction to the R5-R6 region of talin-2.

Together with the single-molecule measurements, these biochemical findings establish that talin-2 contains two spatially separated, mechanically gated AKAP regions, one within R5-R6 and the other associated with the conserved R9 site. Thus, isoform-specific divergence has expanded the repertoire of PKA-binding sites within the talin-2 rod, providing a mechanism through which mechanical forces may differentially regulate the recruitment and spatial organization of PKA signaling.

**Figure 7.**
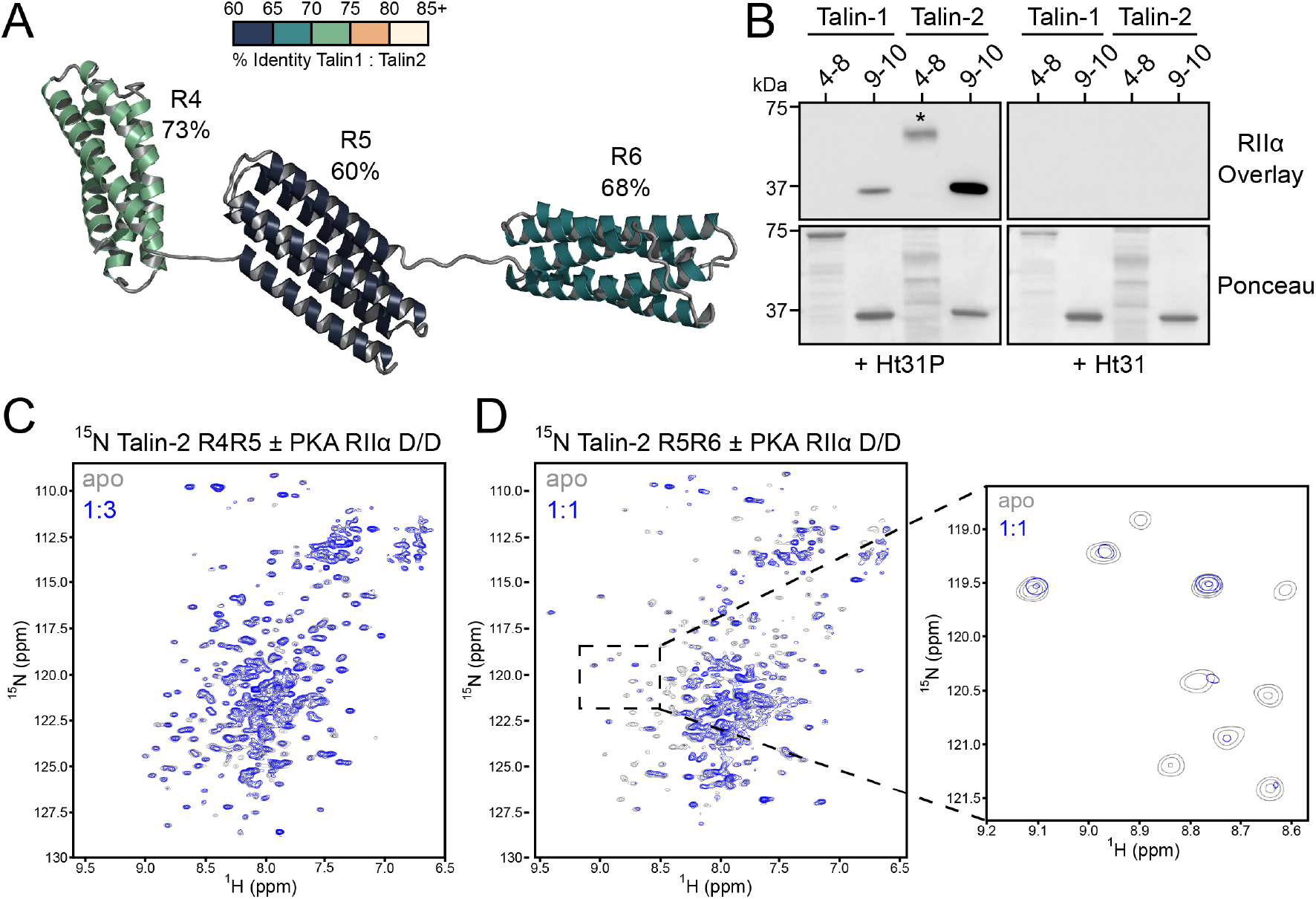
Biochemical identification of an additional PKA-binding region in talin-2 R5-R6. **(A)** Structural representation of talin-2 R4-R6, with individual domains colored according to their percentage sequence identity with talin-1. R5 exhibits the lowest sequence conservation (60%), compared with R4 (73%) and R6 (68%). **(B)** PKA RIIα overlay assays of recombinant talin-1 and talin-2 R4-R8 and R9-R10 fragments in the presence of the inactive control peptide Ht31P (left) or the AKAP-competitive inhibitor Ht31 (right). Talin-2 R4-R8 exhibits an RIIα-binding signal absent from the corresponding talin-1 fragment, while talin-2 R9-R10 produces a substantially stronger signal than talin-1 R9-R10. Both interactions are abolished by Ht31. Ponceau staining confirms comparable protein loading (bottom). The asterisk indicates the RIIα-reactive band corresponding to talin-2 R4-R8. (C,D) Overlay of ¹H,¹⁵N NMR spectra of ¹⁵N-labeled talin-2 R4-R5 **(C)** and R5-R6 **(D)** in the absence (gray) and presence (blue) of unlabeled PKA RIIα dimerization/docking (D/D) domain. Spectra were recorded at talin:RIIα D/D molar ratios of 1:3 for R4-R5 and 1:1 for R5-R6. Addition of RIIα D/D produces little detectable spectral perturbation in R4-R5 but substantial peak broadening and spectral changes in R5-R6, identifying an additional PKA-binding region within talin-2 R5-R6. The expanded region in (D) highlights representative spectral perturbations.

## Discussion

Although talin-1 and talin-2 share a highly conserved domain architecture and broadly similar full-length unfolding behavior, our results show that the two isoforms encode distinct mechanical responses and ligand-binding capabilities. Talin-2 is selectively retuned at key rod segments in ways that redistribute the unfolding hierarchy, alter interdomain coupling, and reshape the force-dependent exposure of cryptic binding sites. These differences are accompanied by changes in the recruitment of both structural and signaling partners, indicating that isoform-specific sequence divergence can diversify mechanotransduction without altering the overall architecture of the talin rod.

The R1-R3 region provides an example of how changes in mechanical stability can tune the force response of the talin scaffold. In talin-1, R3 is the most mechanically labile rod domain, unfolding at forces of approximately 5 pN and acting as a key low-force mechanosensor during adhesion assembly. R3 is also the most mechanically labile domain in talin-2, but is modestly destabilized relative to talin-1, with its unfolding force decreasing from approximately 5 pN to 4 pN. The higher-force unfolding transitions are also redistributed, with some domains unfolding at lower forces and others exhibiting increased mechanical stability in talin-2. Previous studies have demonstrated that the mechanical stability of R3 can be tuned through mutations that alter the packing of its hydrophobic core, shifting the force threshold for unfolding and vinculin recruitment (5, 23). More recently, alternative splicing of TLN1 was shown to introduce a 17-amino-acid insertion between R1 and R2 that alters the mechanical response of this region (24), while targeted disruption of the R1-R2 interface was found to affect stiffness sensing and mechanical homeostasis in mice (25). Previous work by Austen and colleagues also identified isoform-specific differences in talin-mediated mechanical linkages involving the R1-R3 region, supporting the potential functional significance of the mechanical differences observed here (15). Together, these findings highlight the importance of the R1-R3 region as a tunable mechanosensitive module and suggest that differences in its mechanical stability between talin-1 and talin-2 may contribute to isoform-specific responses during force-dependent adhesion reinforcement.

A more striking difference between the isoforms emerges in the central R7-R9 region, where talin-2 exhibits a fundamentally different unfolding hierarchy and a distinct cooperative unfolding pathway. In talin-1, R7 and R8 unfold cooperatively, followed by R9 as a separate, higher-force event. By contrast, talin-2 R7-R9 exhibits two alternative unfolding pathways: one in which R9 unfolds first, followed by the cooperative unfolding of R7-R8, and another in which all three domains unfold together in a single transition. The coexistence of these pathways over overlapping force ranges suggests that the mechanical response of this region is governed not simply by the stability of individual domains, but also by the coupling between neighboring helical bundles. Thus, isoform-specific sequence differences appear to alter the mechanical coupling within R7-R9, introducing a cooperative unfolding pathway in talin-2 that is not observed in talin-1. This altered coupling may have important consequences for the coordinated exposure of ligand-binding sites, providing a mechanism by which mechanical forces could regulate multiple interactions within a single region of the talin scaffold.

The mechanical differences between talin-1 and talin-2 may be explained, at least in part, by isoform-specific changes in the packing of their helical bundles. Talin rod domains are stabilized by tightly packed hydrophobic cores, and previous studies have demonstrated that the introduction of buried polar residues can destabilize individual helical bundles and alter their mechanical stability (7, 25, 26). The best-characterized example is the threonine belt within talin-1 R3, which destabilizes the helical bundle and contributes to its unusually low unfolding force. Substitution of these threonines with hydrophobic residues in the IVVI mutant restores mechanical stability (5). Despite sharing approximately 76% sequence identity, inspection of the talin-1 and talin-2 sequences reveals numerous differences in buried core residues, including substitutions involving serine and threonine. Intriguingly, many of these differences are conserved within each isoform across species, suggesting that they may contribute to the distinct mechanical properties of the two proteins.

Perhaps the most significant consequence of isoform-specific sequence divergence is the emergence of new mechanically regulated ligand-binding sites in talin-2, particularly within the highly divergent R4-R6 region. R5 is the most divergent domain between the two isoforms (Fig. 1A), yet remains highly conserved within each isoform across species, suggesting that these differences have been maintained through evolutionary selection. Our experiments reveal that talin-2 R5 contains a previously unrecognized vinculin-binding site, providing direct evidence that sequence divergence can alter the ligand-binding capabilities of the talin scaffold. Importantly, this site is cryptic in the folded domain and becomes accessible following mechanical unfolding, allowing vinculin to engage and stabilize the unfolded state. Thus, the differences between talin isoforms extend beyond changes in mechanical stability and unfolding hierarchy to include the repertoire of interactions that can be activated by force.

Remarkably, the acquisition of new mechanically regulated interactions within talin-2 R4-R6 is not restricted to vinculin. We also identify an additional PKA-anchoring region within R5-R6, distinct from the previously characterized R9-associated AKAP site shared with talin-1. This indicates that talin-2 contains two spatially separated PKA-anchoring regions, potentially expanding its capacity to organize kinase signaling along the mechanically active scaffold. Interestingly, the R9-associated interaction also appears to differ between the isoforms. In talin-1, robust PKA RIIα binding requires the R9-R10 region, including approximately nine residues from R10, whereas our single-molecule experiments demonstrate PKA-dependent suppression of refolding within talin-2 R7-R9, even in the absence of these additional residues. The stronger RIIα overlay signal observed for talin-2 R9-R10 suggests enhanced RIIα binding, while the PKA-dependent suppression of R7-R9 refolding raises the possibility that the talin-2 AKAP interaction is less dependent on the structural contributions of R10. Together, these findings establish that talin isoforms differ in both the spatial organization and mechanical regulation of PKA-anchoring sites, providing distinct mechanisms for coupling mechanical forces to local phosphorylation signaling within adhesions.

Talin-2 is the predominant isoform in the brain (27) and is also enriched in large, mature adhesions (28, 29) and mechanically demanding structures such as costameres (30) and myotendinous junctions, suggesting that its altered mechanical properties and expanded ligand-binding capabilities may be important across diverse cellular environments. Rather than functioning as interchangeable scaffolds, the two isoforms appear to encode distinct responses to mechanical force, with differences in domain stability, unfolding cooperativity, and the recruitment of structural and signaling partners. These findings also have implications for how mechanical information is encoded within the talin scaffold. The helical bundles of the talin rod function as mechanically gated binary switches, transitioning between folded and unfolded states to expose or conceal ligand-binding sites. By changing the stability and coupling of these switches, isoform-specific sequence differences can generate distinct patterns of domain activation under comparable mechanical inputs. Within the MeshCODE framework (3, 31, 32), these patterns represent different mechanically encoded states of the talin scaffold, raising the possibility that talin-1 and talin-2 provide cells with distinct molecular mechanisms for interpreting and responding to their mechanical environment.

The structural basis of the distinct cooperative unfolding behavior observed in talin-2 R7-R9 remains to be established, particularly whether this behavior arises from differences in interdomain interfaces, helical bundle packing, or force propagation between adjacent domains. Similarly, further work will be required to define the molecular basis of the newly identified vinculin- and PKA-binding interactions and determine how these differences influence ligand recruitment and signaling in living cells. Although talin-1 and talin-2 share approximately 76% sequence identity (11), systematically interrogating the remaining 24% sequence divergence will be important for identifying the substitutions responsible for differences in mechanical stability and ligand recruitment, as well as isoform-specific post-translational modification sites that may further diversify their signaling properties.

The altered mechanical responses of the two talin isoforms, together with the convergence of additional vinculin- and PKA-binding activities within talin-2 R4-R6, highlight how isoform-specific sequence divergence can reshape the mechanical and biochemical properties of the talin scaffold, promoting the assembly of distinct structural and signaling complexes in a force-dependent manner. These findings reveal how highly conserved mechanosensitive proteins can evolve specialized mechanical and signaling functions within a shared structural framework, providing a molecular basis for understanding the nonredundant roles of talin isoforms in cellular mechanotransduction.

## Materials and Methods

### Cloning, expression and purification of recombinant proteins

Plasmids encoding the R4, R5, R6 and R4-R8 rod domains of talin-2 were purchased as synthetic genes cloned into pET151/D-TOPO (Life Technologies). The PKA RIIα docking/dimerization (D/D) domain (residues 1-50) was expressed from a pET28 construct.

DNA sequences encoding the talin-2 R4-R5 and R5-R6 domains were amplified from the pET151/D-TOPO R4-R8 construct using primers containing 5′ extensions complementary to the multiple cloning site (MCS) of pET47b, and subcloned using ligation-independent cloning (LIC). The 3′ to 5′ exonuclease activity of T4 DNA polymerase was used to generate complementary 5′ overhangs on the PCR products and XmaI/SacI-digested vector. The PCR products and vector were annealed at room temperature, and the resulting DNA was used to transform XL2-Blue competent cells (Agilent Technologies). Individual transformants were cultured in LB medium supplemented with 50 μg mL⁻¹ kanamycin, and plasmid DNA was purified using a Monarch Plasmid Miniprep Kit (NEB). Plasmid sequences were verified by Sanger sequencing.

Mechanically stretchable talin-2 constructs corresponding to the full rod (R1-R13) and subfragments (R1-R3, R4-R6, R7-R9 and R9-R12) were expressed and purified as described previously (8). Briefly, recombinant proteins were purified via an N-terminal GST tag using glutathione Sepharose resin (GE Healthcare), followed by tag removal through TEV protease cleavage.

His-tagged talin-2 rod fragments were expressed in *Escherichia coli* BL21(DE3)pLysS cells (Novagen) grown in LB medium or 2M9 minimal medium supplemented with ¹⁵NH₄Cl for isotopically labeled proteins. Expression was induced at an OD₆₀₀ of 0.7 with 625 µM IPTG for 16 h at 18 °C. Proteins were purified by nickel-affinity chromatography using a HisTrap HP column (Cytiva), followed by anion- or cation-exchange chromatography using HiTrap Q HP or HiTrap SP HP columns (Cytiva), respectively. Purified proteins were buffer-exchanged into SEC buffer (50 mM Tris-HCl, pH 7.4, 150 mM NaCl, 1 mM TCEP) or NMR buffer (20 mM sodium phosphate, pH 6.5, 50 mM NaCl), concentrated and stored at −80 °C.

The vinculin D1 domain (Vd1) was cloned into the pET151 expression vector as previously reported (33). His-tagged Vd1 was purified using immobilized metal affinity chromatography, followed by anion-exchange chromatography as described previously (34).

The PKA RIIα docking/dimerization (D/D) domain was expressed and purified as described previously (19). Protein concentrations were determined spectrophotometrically at 280 nm using calculated extinction coefficients. For talin-2 R4 and R6, which lack appreciable intrinsic absorbance at 280 nm, the tyrosine-containing TEV cleavage site was retained to enable spectrophotometric determination of protein concentration.

### RII overlay

Recombinant proteins (1.5 µg) were separated by SDS-PAGE and transferred to nitrocellulose membranes. Membranes were blocked with 10% nonfat dry milk (NFDM) in TBS-T for 1 h and incubated with V5-tagged full-length PKA RIIα D/D domain (0.73 µg/mL) in 10% NFDM for 12 h. Where indicated, Ht31 or Ht31P peptides were included at 1 µM. Following washing with TBS-T, membranes were incubated with V5-HRP (BioLegend, 1:4,000 in 10% NFDM) for 12 h, washed with TBS-T, and developed by enhanced chemiluminescence.

### Size-exclusion chromatography (SEC)

Analytical SEC was performed using a Superdex 75 10/300 Increase (Cytiva) equilibrated in 50 mM Tris-HCl (pH 7.4), 150 mM NaCl and 1 mM TCEP. Talin-2 rod domains were incubated with Vd1 at a 1:1 molar ratio for 16 h at 42 °C before analysis. Samples containing 100 μM of each protein were separated at a flow rate of 750 μL min⁻¹, with UV absorbance monitored at 214, 260 and 280 nm. Fractions (250 μL) were collected and analyzed by SDS-PAGE.

### Nuclear magnetic resonance (NMR) spectroscopy

¹⁵N-labeled protein samples were prepared at 100 μM in 20 mM sodium phosphate, pH 6.5, 50 mM NaCl, 1 mM DTT and 5% (v/v) D₂O. ¹H and ¹H,¹⁵N TROSY spectra were collected at 298 K using a Bruker Avance 700 MHz spectrometer equipped with a cryogenic probe at the University of Liverpool High-Field NMR Facility. Unlabeled PKA RIIα D/D domain was added to ¹⁵N-labeled talin-2 R4-R5 or R5-R6 at molar ratios of 1:1 and 1:3 to assess binding-induced spectral changes. Data analysis was performed using TopSpin 4.5.0 and CCPNMR Analysis 3.2.2 (35).

### Single-molecule manipulation

Single-molecule experiments were performed as described previously (8) using a custom-built high-force magnetic tweezers instrument capable of applying forces up to 100 pN with ∼1 nm spatial resolution (36, 37). Talin-2 fragments (R1-R3, R4-R6, R7-R9, and R9-R12) were immobilized on glass coverslips within a laminar flow chamber via HaloTag-Halo ligand coupling at the C-terminus and tethered to 3-µm paramagnetic beads through an N-terminal biotin-streptavidin linkage.

Measurements were performed in PBS (pH 7.4) supplemented with 2 mM TCEP. To minimize force artifacts during introduction of Vd1 or PKA, a buffer-isolation membrane well array was used to separate reagent exchange from the measurement chamber (38). Force spectroscopy measurements were conducted at constant loading rates over a narrow range of 2-4 pN s⁻¹ as specified in the corresponding figure legends.

Unless otherwise stated, each dataset represents measurements from multiple independent tethers and several hundred unfolding events.

Force calibration was performed using a standard calibration curve (36), requiring a single reference point relating magnet-bead distance to applied force. The DNA overstretching transition at ∼65 pN (39) was used as the calibration reference. The estimated uncertainty in force determination was approximately 5%.

## Acknowledgements

J. Y. was supported by the Singapore Ministry of Education Academic Research Fund Tier 2 (MOE-T2EP50123-0008 to J.Y.) and Tier 3 (MOE-MOET32021-0003 to J.Y.), the National Research Foundation Singapore through the Mechanobiology Institute Mid-Sized Grant (NRF-MSG-2023-0001 to J.Y.), and the Research Grant from HFSP (Ref.-No: RGP025/2024). B.T.G. was supported by BBSRC grants (BB/N007336/1 and BB/S007245/1), an HFSP grant (Ref.-No: RGP025/2024), and a British Heart Foundation grant (SP/F/23/150045). D.M.C. was supported by the Lyons-Jones Scholarship (UKG10010), funded through an endowment to the University of Liverpool.

## Data Availability

All data supporting the findings of this study are available within the article. Additional data are available from the corresponding author upon reasonable request.

